# Differential Responses of Dynamic and Entropic Aging Factors to Longevity Interventions

**DOI:** 10.1101/2024.02.25.581928

**Authors:** Kristina Perevoshchikova, Peter O. Fedichev

## Abstract

Aging across most species, including mice and humans, is characterized by an exponential acceleration of mortality rates. In search for the molecular basis of this phenomenon, we analyzed DNA methylation (DNAm) changes in aging mice. Utilizing principal component analysis (PCA) on DNAm profiles, we identified a primary aging signature with an exponential age dependency, closely reflecting the Gompertz law’s description of mortality acceleration. This signature is the manifestation of the dynamic instability in the organism’s state that drives the aging process in mice. It aligns closely with regression-based aging clocks and responds to interventions such as caloric restriction and parabiosis. Additionally, we identified a linear DNAm signature, indicative of a global demethylation level. Through single-cell DNAm (scDNAm) data from aging animals, we demonstrate that this signature captures the exponential expansion of the state space volume spanned by individual cells within an aging organism, and thus quantifying linearly increasing configuration entropy, likely an irreversible process. Consistent with this interpretation, we found that neither caloric restriction (CR) nor parabiosis significantly impacts the entropic feature, reinforcing its link to irreversible damage.

## I. INTRODUCTION

Aging in most species manifests itself as an exponential acceleration of mortality. Practical biotechnology relies on manipulation of biological systems at the molecular and cellular levels and necessitates an understanding of how such exponential dynamics are mirrored by molecular changes, including target molecules concentrations and DNA methylation (DNAm) levels. Concurrently, significant efforts are directed towards developing “aging clocks” or “biomarkers of aging” [1, 2], typically derived using supervised learning techniques. These methods involve fitting multidimensional signals, such as DNAm profiles or blood markers, to chronological age [3, 4] or mortality risk [5, 6]. However, overcoming the curse of dimensionality in these analyses necessitates aggressive regularization. For instance, the best-known Horvath DNAm clock [3] was derived from ultra-high dimensional data using Elastic Net, a combination of L1 and L2 regularization [7].

The heavy reliance on regularization presents multiple challenges. Primarily, achieving a unique solution is challenging, as heavy regularization tends to select a number of features approximately equal to the sample size, based on their correlation to the target phenotype. Consequently, the feature sets identified through regression may lack direct biological relevance, especially once the number of features substantially exceeds the sample size. As a result, our “biological clocks” may more accurately reflect preconceived notions about the aging process, such as the linear dependence of key aging features on chronological age. These predictors then require additional statistical models, such as log-linear proportional hazards models [6], to associate the identified linear features with the observed exponential increase in mortality.

An effective alternative is the use of unsupervised approaches, such as principal component analysis (PCA, see, e.g. [8]), non-negative matrix factorization (see example of NMF applied to DNAm data [4]), or modern AI/ML systems designed for longitudinal biomedical data analysis [9, 10]. These methods aim to represent the data using a limited number of variables while maintaining controlled accuracy, either at a single time point or across aging trajectories. The key advantage of these models is their minimal underlying assumptions, allowing for a more authentic revelation of the dynamics of age-related features.

In this Letter, we followed [11] and utilized PCA to obtain a semi-quantitative understanding of the dynamics of methylation profiles associated with aging, as well as their response to lifelong and transient interventions in mice. This was done using bulk DNA methylation (DNAm) data from [4] and newly available single cell DNAm (scDNAm) data from [12]. We produced evidence suggesting that the leading aging signature in the bulk DNAm data from mice characterizes the dynamics of a large cluster of tightly-synchronized age-dependent features exhibiting an exponential dependence on age with the exponent matching the exponent of the Gompertz mortality law. The variance of the the feature increased exponentially within the population, albeit at twice the rate. Such a pattern of exponential growth in both mean and variance is indicative of stochastic instability of the organism state, and hence we identify DNAm-PC1 with the order parameter characterizing the development of the instability, the dynamic frailty index (dFI, see [9]). Our DNAm estimate of dFI correlates with and is enriched with DNAm sites selected by aging clocks (trained to predict the chronological age of animals [4, 13]) commonly used in the experiment in mice and responded to short-term (parabiosis) and long-term (caloric restriction, CR) treatments.

The next leading DNAm signature identified in our study is linear over time, reflecting a global trend of demethylation of mouse DNA in the course of aging process. Through the analysis of scDNAm data, we demonstrate that most of these DNAm features alter their methylation states independently. This behavior supports our ealier hypothesis [11] suggesting that linear features in DNAm measurements track an exponential expansion of the volume of configuration state space volume occupied by single cells in the aging organism, a likely irreversible process. Thus, the linear aging signature may represent irreversible damage measured by increasing the configuration entropy. Consistent with these expectations, analysis of publicly available data revealed that neither caloric restriction (CR) nor parabiosis can significantly reduce or affect the rate of change of this linear feature, reinforcing its association with irreversible entropy accumulation.

Finally, we draw comparisons between the leading aging signatures in mice and humans. Despite the observed qualitative differences in aging dynamics between the two species, our analysis suggests that aging in both can be quantitatively described by the same overarching model using a physics-informed aging model from [9, 11, 14, 15]. This alignment, along with validation of the entropic nature of the leading aging signature, opens opportunities for the discovery and testing of experimental interventions that could significantly affect human lifespan using short-lived species like rodents as models.

## II. RESULTS

### A. Aging and DNAm in mice

The GSE80672 data set comprises the bulk DNAm levels from two tissues of from C57BL/6 mice. In this study,we focused exclusively on blood samples (the same data was used in the development of Petkovich’s DNAm clock, which is often used in mouse experiments [4]). To discern clusters of correlated features in the DNAm signal and understand the character of their variation with age, we conducted a Principal Component Analysis (PCA) after filtering out sites with the weakest correlation to chronological age (details in Materials and Methods S.I A).

Our findings are presented in Figs. 1. For each sample, PCA yields scores, henceforth referred to as DNAm-PC1 and DNAm-PC2, representing the common factors behind the fluctuations of two largest clusters of DNAm sites. We applied an exponential fit to individual PCA scores across all samples, revealing that DNAm-PC1 increases exponentially with age, with an estimated exponent of *α* = 0.17 per month (see Fig. 1(a)). The exponent is not small, *αt*_*ls*_ ≈ 5 ≫ 1, where *t*_*ls*_ ≈ 28 months is the average life expectancy of the C57BL/6 mice [16].

**FIG. 1.**
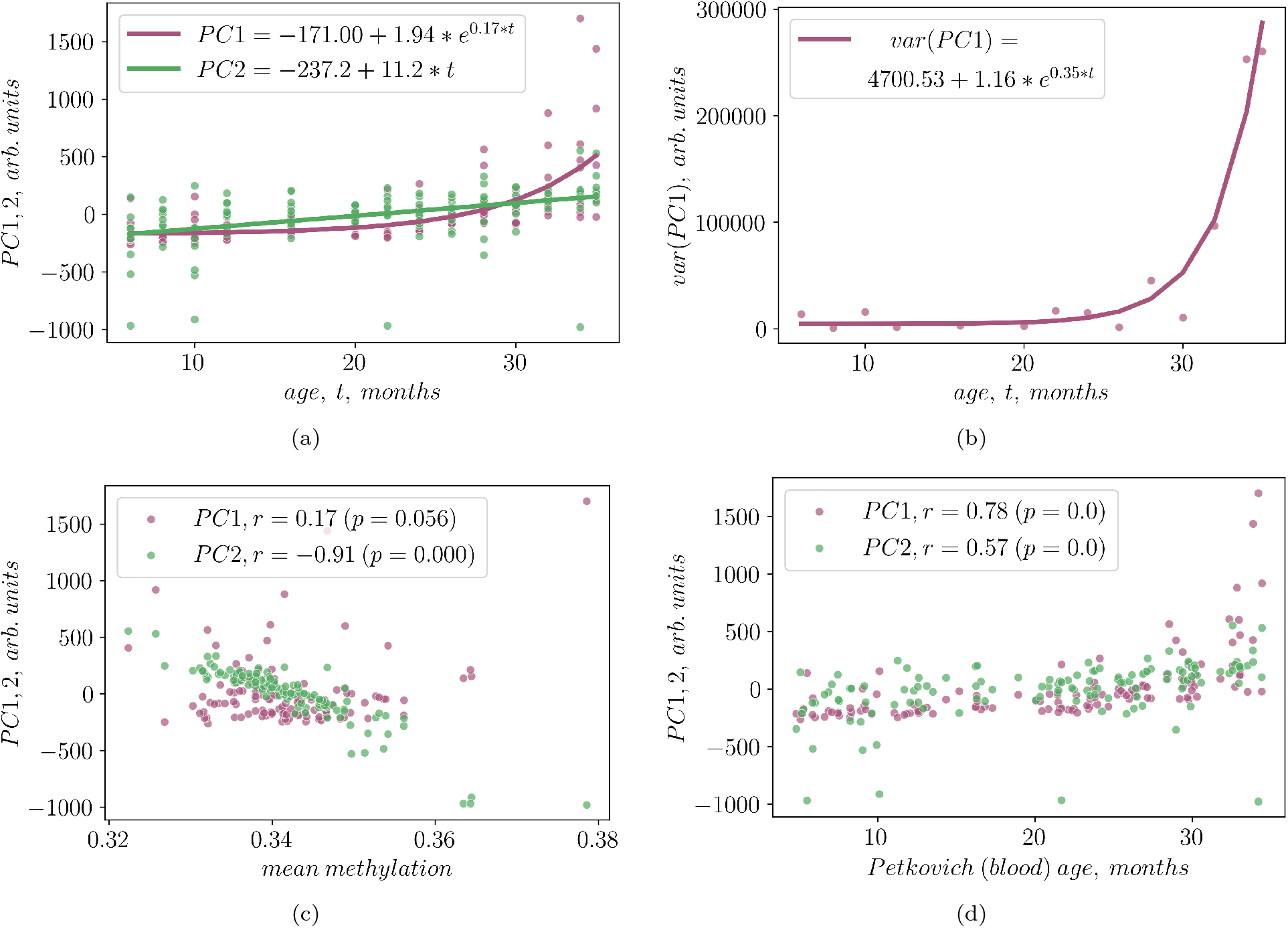
Analysis of Aging Dynamics of DNA Methylation (DNAm) States Using PCA. Panel (1(a)): Illustrates the leading aging signatures in bulk DNAm, revealing two clusters of DNAm features with exponential and linear dependencies on chronological age. Panel (1(b)): Demonstrates that the variance of the exponential DNAm-PC1 signature exhibits an exponential increase with age. Panel (1(c)): Shows the correlation between the linear DNAm feature and the mean DNAm level. Panel (1(d)): Highlights the correlations betweem the DNAm signatures with the aging clock developed in the same dataset and described in [4].

In addition to the exponential growth observed in the mean of DNAm-PC1, we found a similar exponential increase in its variance with age. This increase was approximately twice the rate of the mean DNAm-PC1 score, as shown in Fig. 1(b). Such a pattern of exponential growth in both mean and variance is indicative of stochastic instability of the organism state, and hence we identify DNAm-PC1 with the order parameter characterizing the development of the instability, the dynamic frailty index (dFI) in mice [9].

On the contrary, DNAm-PC2 exhibited a distinctly different pattern. Its relationship with age is best described as a linear function (Pearson correlation coefficient of *r* = 0.42 (*p* = 1.04 · 10^*−*6^), see Fig. 1(a)) with no evidence for increasing variance over time.

The DNAm-PC2 score has a straightforward interpretation: it correlates with the overall average methylation level in our samples (Pearson’s correlation coefficient of *r* = *−*0.91 (*p <* 10^*−*3^), see the green dots in Fig. 1(c)). Conversely, DNAm-PC1 does not show a notable correlation with average methylation levels, as indicated by the magenta dots in the same figure (Pearson’s correlation coefficient of *r* = 0.17 (*p* = 0.056), see the green dots in Fig. 1(c)).

Finally, we investigated the relationship between DNAm-PC1 and DNAm-PC2 scores, on the one hand, and the DNAm clock developed using this dataset [4], on the other (see Fig. **??**). Our analysis revealed significant correlations for both DNA-PC scores. This is not surprising, since both PC scores correlate to the common factor – the chronological age. However, the exponential feature (DNAm-PC1) more closely mirrored the aging dynamics captured by the methylation clock from [4] than the linear feature (DNAm-PC2).

PCA transformation produced the PC scores and the corresponding loading vectors. The loading vector components have the meaning of the degree of participation of each DNAm site in the dynamics of the corresponding cluster of DNAm features. The probability distributions of the components of the loading vector (also known as participation indices) for DNAm-PC1 and DNAm-PC2 are represented in magenta and green colors, respectively, in Fig. 2(a).

**FIG. 2.**
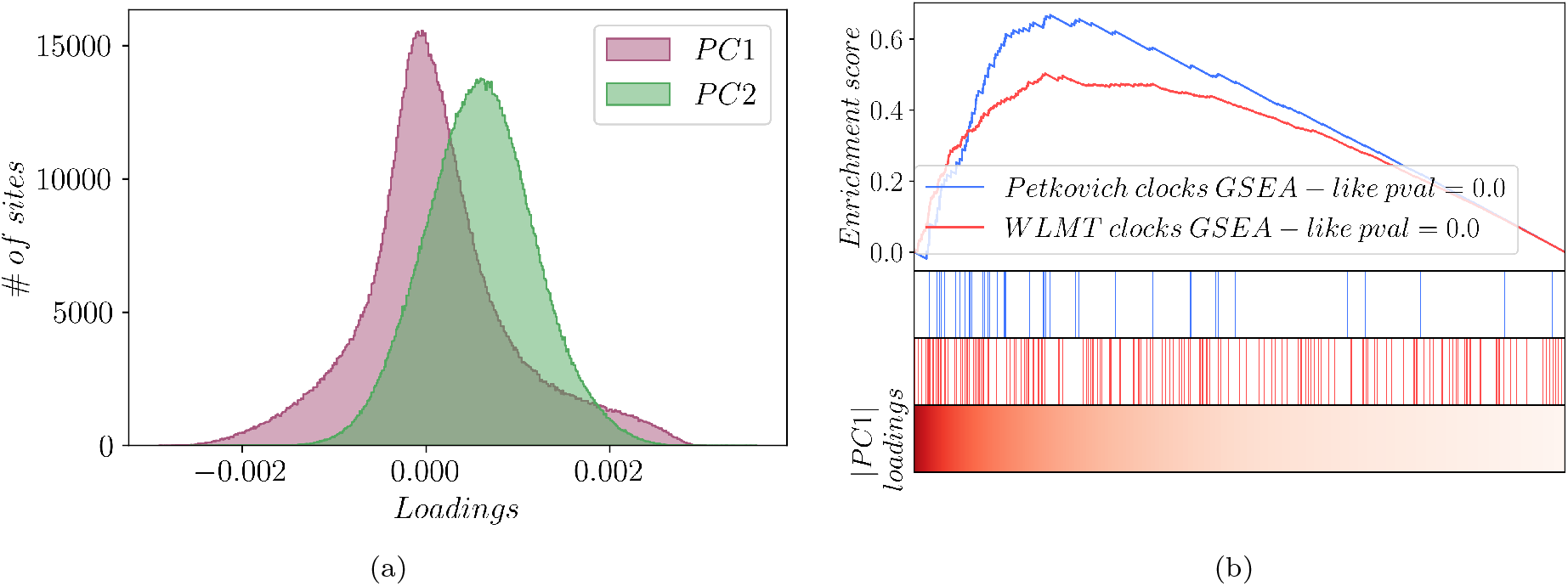
Properties of the loading vectors corresponding to the leading exponential (DNAm-PC1) and linear (DNAm-PC2) features. (2(a)) Loading vectors (participation coefficients) of the exponential and the linear aging signatures show qualitatively different distributions; (2(b)) Leading (max-absolute value) components of the DNAm-PC1 loading vector are enriched with DNAm sites selected for biological age determination in Petkovich [4] and WLMT [13] clocks.

We note, that due to the limited availability of DNAm samples, coupled with the ultra-high dimensionality of the signal, makes it challenging to precisely delineate the features contributing to the DNAm-PC1 and -PC2 signatures. Furthermore, with an increase in the number of samples, matrix-factorization-based methods like PCA are likely to reveal additional processes beyond aging reflected in the data. These higher-order contributions would correspond to specific biological processes.

The distribution of the loading vector components for the exponential feature, DNAm-PC1, displays heavy tails, indicating the presence of sites significantly associated with this process. We performed a gene set enrichment analysis (GSEA) for these relevant sites to identify genes likely modulated by DNAm-PC1 activation and enriching developmental and signaling pathways (30 pathways passing FDR criterion at *q <* 1 · 10^*−*4^ are presented in Supplementary Table 1). These observations match earlier findings in the same dataset [17].

In contrast, the components of the DNAm-PC2 loading vector corresponding to the linear feature show a distribution shifted away from zero, implying a non-localized activation and hence representing the change in the average methylation level (in line with Fig. 1(c)). Due to the global nature of this feature, there was no sense in conducting GSEA.

### B. DNAm-PC scores and aging clocks

The exponential signature DNAm-PC1 is significantly enriched with DNAm sites commonly selected by supervised aging clocks. We organized DNAm sites according to the absolute values of PC1 loading vector components in ascending order, as illustrated in the bottom panel of Fig. 2(b). The sites employed in the regularized regression for the construction of the Petkovich[4] and WLMT [13] aging clocks are marked, respectively, in blue and red.

Enrichment analysis showed that sites chosen by both aging clock regression models likely correspond to the leading components of the DNAm-PC1 loading vector, both for Petkovich (Minimal Hypergeometric (mHG) *p* = 4.74 *·* 10^*−*9^) and for WLMT (*p* = 7.55 *·* 10^*−*14^) clocks.

### C. Aging and configuration entropy - evidence from single cell methylation

The most plausible interpretation of the observed linear aging trajectories shared by a wide array of DNAm features is that the dynamics of DNAm-PC2 is governed by numerous independent alterations in methylation states. This theory, originally proposed in [11] concerning human DNAm data, finds indirect support through the observed correlation between DNAm-PC2 and average methylation in mice, as illustrated in Fig. 1(c).

To validate this interpretation on the level of individual cells, we analyzed single-cell DNAm (scDNAm) data from [12], calculating mutual information (MI) between methylation sites to assess codependence (detailed in Materials and Methods). This nonnegative measure is exactly zero if the sites change their states independently(see, e.g., [18]).

Given the sparsity of scDNAm data, we relied on one more level of averaging and computed the average MI for each DNAm site, arranged them by increasing mean MI (see the bottom panel in Fig. 3), and marked sites integral to the DNAm-PC1 loading vector.

**FIG. 3.**
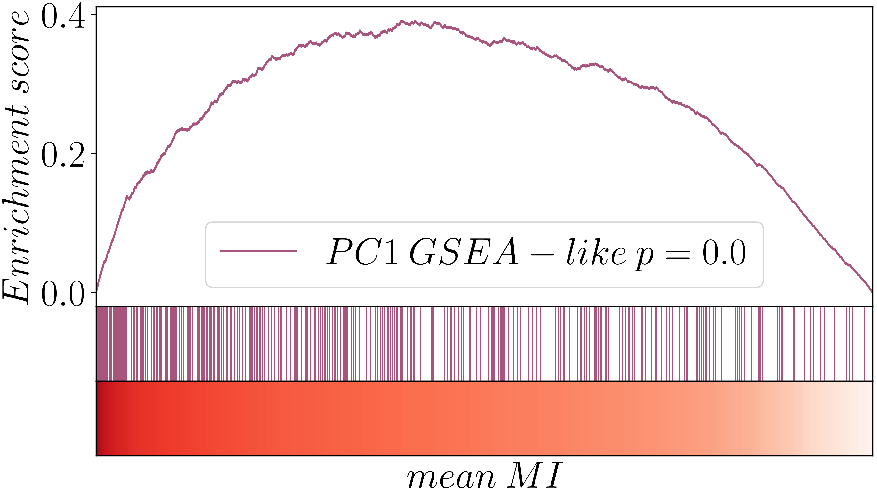
The distribution of the leading DNAm-PC1 loading vector components (bars) corresponding to DNAm sites ordered by the average mutual information (MI, the color bar in the bottom panel). High-average MI sites change their states in a correlated manner and more likely to belong to DNAm-PC1 signature (mHG *p* = 8.215 *·* 10^*−*9^)

The distribution of average MI for DNAm-PC1’s leading components is left-shifted (mHG *p* = 8.215 · 10^*−*9^), implying concurrent methylation changes of DNAm sites associated with dynamics of DNAm-PC1. This finding, together with the exponential dependence of mean and variance on age, reinforces the link between the DNAm-PC1 score and the order parameter characterizing the dynamic instability of the organism state - the dynamic frailty index, dFI [9]. Therefore, we propose using the DNAm-PC1/dFI to encapsulate the dynamic aspects of aging.

Conversely, DNAm-PC2-associated sites show a lower average MI, suggesting that most of DNAm states change independently. Therefore, the volume in the configuration space occupied by individual cellular states within an aging organism grows exponentially with age, and hence configuration entropy scales linearly with time. Following [11], we identify this process as DNAm-PC2 with thermodynamic biological age (tBA, introduced in [11]), henceforth referred to as DNAm-PC2 / tBA, capturing the entropic component of aging.

### D. Effects of interventions on aging signatures

We investigated the response of linear(entropic) and exponential(dynamic) DNAm signatures to both lifelong and short-term interventions (see Figs. 4). Caloric restriction (CR), recognized for its lifespan extension effects in mice, was examined using DNAm data from [4]. Our analysis also showed that CR markedly decreased the growth rate of the dynamic signature, DNAm-PC1/dFI (Fig. 4(a), *p* = 0.0, two-sided t-test). However, CR did not significantly alter the rate of entropy production, as measured by the change in the slope of age dependence of the DNAm-PC2/tBA score (Fig. 4(b), *p* = 0.34, two-sided t-test).

**FIG. 4.**
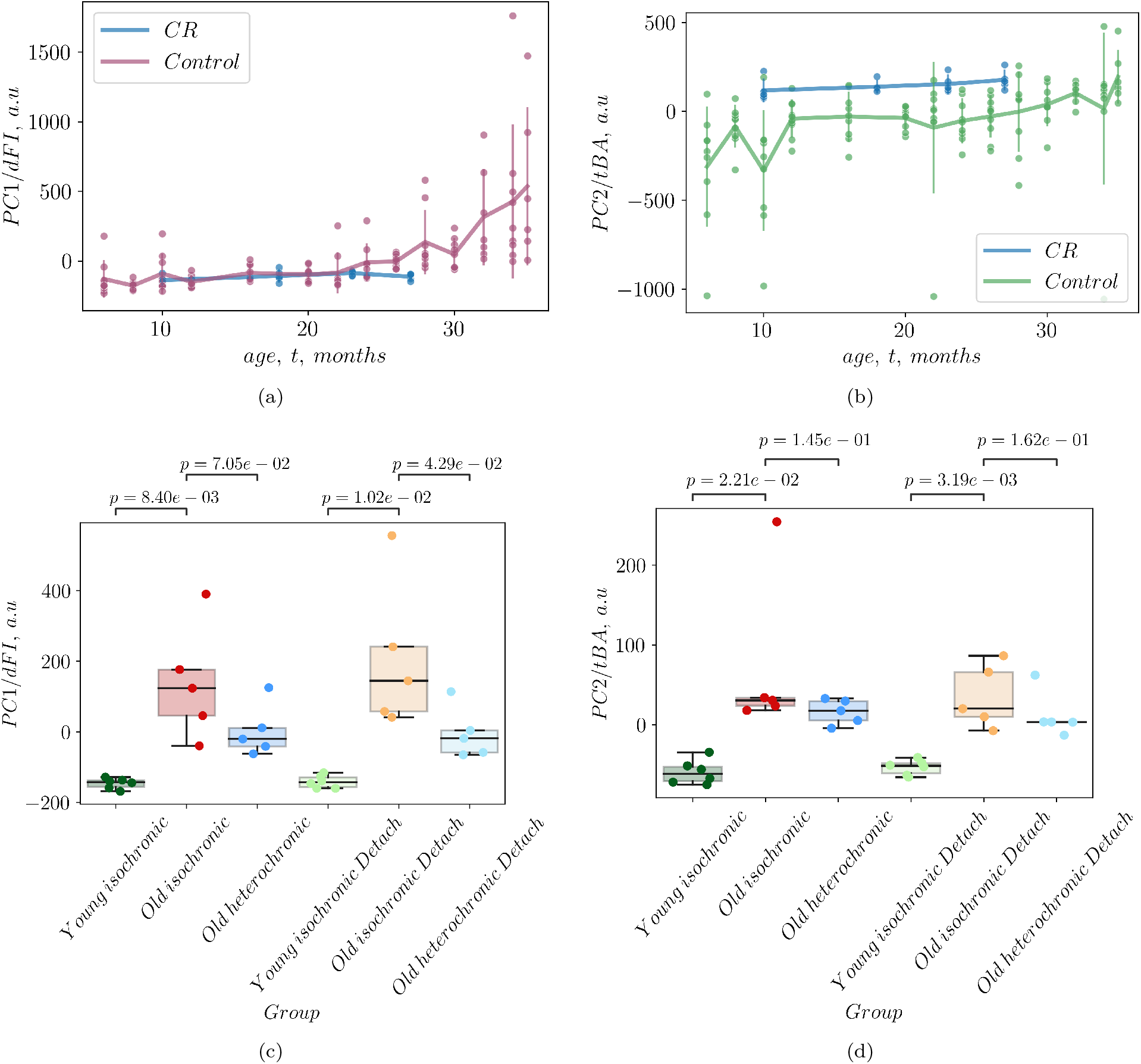
Effects of life-extending interventions, caloric restriction (CR, top), and parabiosis (bottom), on dynamic (DNAm-PC1/dFI, left) and entropic (DNAm-PC2/tBA, right) aging components, respectively.

We observed that DNAm-PC2 levels are higher in samples from CR animals compared to controls (blue relative to green dots in Fig. 4(b), *p* = 0.025, two-sided t-test). However, we caution against drawing extensive conclusions from this observation, as life-long CR might influence higher-order adaptive processes not directly related to aging, which could result in systematic shifts in DNAm. Accurate estimation of DNAm-PC scores characterizing DNAm profiles with lesser variance would require significantly larger sample sizes. Nevertheless, this uncertainty does not impact our slope estimates and hence alter our findings on the effects of CR on aging rates as measured by tBA.

In the case of heterochronic parabiosis, known to extend lifespan in mice, the application of our PCA-based model to DNAm data from [19] revealed that the dynamic/exponential (DNAm-PC1/dFI) and the linear/entropic (DNAm-PC2/tBA) signatures were markedly elevated in older mice relative to younger counterparts. This observation, leveraging a dataset not involved in our PCA model training, externally validates our PCA-identified aging signatures.

In parabiosis experiments, the dynamic DNAm-PC1/dFI signature was significantly reduced (*p* = 7.05 · 10^*−*2^, one-sided t-test) during the procedure and the effect persisted for 2 months after detachment (*p* = 4.29 · 10^*−*2^, one-sided t-test, see Fig. 4(c)). In contrast, the entropic signature, DNAm-PC2/tBA, remained unchanged immediately after parabiosis and two months later (Fig. 4(d)). Isochronic parabiosis did not show an impact on either of the aging signatures.

## III. DISCUSSION

In this study, we built on our previous work [11] and employed unsupervised learning techniques, specifically principal component analysis (PCA), to uncover the leading DNAm aging signatures in mice. The leading signature, the first principal component DNAm-PC1, demonstrated an exponential correlation with chronological age, with an exponent *α* ≈ 0.15 per month. This exponent closely mirrors the Gompertz mortality law exponent *α*_*G*_ = 0.16–0.24 per month, as detailed in our parameter estimates in [9], using mortality data from healthy controls in [20]. The variance of this exponential feature doubles this rate.

The results could be made sense with the help of the theoretical framework [9, 14], suggesting that the exponential increase in DNAm-PC1, along with its stochastic broadening over time, implies that aging in mice stems from the dynamic instability of the organism’s state. Furthermore, the direct link between DNAm-PC1 scores and the macroscopic trait of accelerating mortality frames aging as an emergent property, measurable by the order parameter—the dynamic frailty index (dFI).

The dFI acts as a common factor for fluctuations in a large cluster of correlated features throughout the organism, rather than any specific subsystem. Hence in no way we assume that DNAm is somehow the most important factor driving the aging process. Rather, DNAm drift—alongside other microscopic features are manifestations of aging sharing universal (system’s independent) properties.

For species with a sufficiently long lifespan, *αt*_*ls*_ *≳* 1, the effects of stochastic forces on the aging trajectory are small [14], and the order parameter, dFI, closely matches the leading first PC score. Therefore, DNAm-PC1 can effectively represent an estimate of dFI from a specific dataset (hence the notation DNAm-PC1/dFI).

On a more practical level, the association of aging with the dynamic instability of the organism’s state suggests that aging in mice is largely reversible [9]. Therefore, interventions targeting dFI should have lasting effects, even after short-term treatments, with the effects of repeated treatments possibly accumulating over time.

Consistent with the theoretical understanding, we observed that dynamic aging signatures are responsive to short-term interventions like parabiosis (as shown here, see Fig.4(c)) and rapamycin (as reported in [9]), as well as long-term interventions such as caloric restriction (CR, discussed here, see Fig.4(a)) and the contrasting impacts of a high-fat diet [9].

PCA analysis of DNAm indicates that the exponential instability of an organism’s state constitutes the main driver of the aging process in aging mice. The next leading DNAm signature of aging in bulk DNAm data captures progressive demethylation with age, a well-documented phenomenon, highlighted in relation to the bulk DNAm dataset utilized in this study through non-negative matrix factorization (NMF), an approach similar in spirit to PCA [4].

The emergence of features linearly correlated with chronological age appears to be universal: the predominant DNAm-PC signature of bulk DNAm from human whole blood samples is also linear with age, exhibiting linearly increasing variance [11]. Based on these statistical properties, we proposed that the linear trend in average methylation suggests DNAm alterations result from a vast array of independent events - configuration transitions occurring at nearly age-independent rates. This process is very distinct from the single-factor model underlying the exponential feature, which implies highly correlated changes among the involved sites.

To corroborate this interpretation, we obtained single-cell DNAm (scDNAm) data and investigated pairwise mutual information (MI) between DNAm sites contributing to the DNAm-PC1 and PC2 signatures. This non-negative metric is zero when two sites alter states independently [18]. In line with our theoretical predictions, sites involved in the linear DNAm-PC2 signature display the lowest mean pairwise MI values, whereas those linked to the exponential DNAm-PC1 signature exhibit higher mean MIs (Fig. 3).

Thus, the linear DNAm signature likely reflects the exponential expansion of the configuration space volume occupied by single-cell states, accompanied by a linear increase in entropy, suggesting an irreversible process. A linear increase in configuration entropy suggests a steady growth in the disorder of the system or the information needed to specify and, if needed, to control the microscopic (that is methylation in our example) state of the organism. Following [11], we advocate for the application of the linear DNAm signature as a measure of thermodynamic biological age (DNAm-PC2/tBA in this study, or simply tBA).

The link between the linear feature, DNAm-PC2 in mice, and configuration entropy might be empirically strengthened by examining the impact (or lack thereof) of life-extending interventions on tBA. We observed that caloric restriction did not significantly the linear slope of age-dependent DNAm-PC2/tBA increase – the entropy production estimate. Furthermore, the linear DNAm aging signature remained unaltered immediately following parabiosis and two months later after the detachment.

The apparent irreversibility of the linear aging signature stems from its nature as a cumulative effect of individually infrequent events tied to transitions between metastable states. Each transition, a form of damage, is safeguarded by substantial activation barriers, ensuring low frequencies for both direct and reverse transitions. This condition renders reversing these changes challenging through weak perturbations. Stronger interventions might unpredictably influence other high-barrier states, indicating that the linear aging signature may resist reversal through feasible interventions. Essentially, this linear signature encapsulates ‘entropic’ and ‘irreducible’ damage, marked by transitions in configuration and regulatory errors with extremely long lifetimes.

The effects of noise on aging and noise-based biomarkers of aging were discussed in [11, 21, 22]. Our study delineates aging signatures as reversible (dynamic) and irreversible (entropic) with markedly different fluctuation properties. If our interpretation is correct, the biological data must exhibit two distinct types of stochastic variance. First, fluctuations in degrees of freedom linked to the order parameter, such as those found in dynamic frailty index (dFI)-like features, should vary more significantly across different organisms while remaining highly synchronized within the same individual (high-dFI states are characterized by low entropy since variance in dFI involves changes in states with high mutual information (MI); this synchronicity among aging hallmarks is empirically observed in experiments where drugs acting against a hallmark of aging (an age-dependent factor) act on others as well). We demonstrated, that fluctuations of dFI can grow exponentially and hyperbolically with age in dynamically unstable and ustable situations, respectively exepmlified by mice (here and in [9]) and humans [11, 15].

Second, features related to the thermodynamic biological age (tBA) are influenced by processes that remain largely independent within a single organism, indicated by low pairwise MI among the involved states. The variability in these features is expected to be consistent across organisms of the same age or even cells within the same organism. We anticipate that future high-quality scDNAm data will validate this expectation.

The coregulated and stochastic clusters of methylation sites, the analogues of the the exponential/dynamic and linear/entropic features from this work, were also described in [23]. The conclusion was that epigenetic aging involves coregulated changes, but it is dominated by the stochastic component. Here, we argue that in sufficiently long-lived species, such as mice with *α*_*g*_ *≳ t*_*ls*_, the exponential instability dominates the dynamics of the organism state and contributes most to all-cause mortality and hence regulates longevity [9, 14].

The dominant aging signature is exponential (dynamic) in mice and linear (entropic) in humans. The next leading aging signature in human bulk DNAm is hyperbolic, not exponential [11]. The reversed sequence of these signatures in PCA analysis and their distinct functional forms are likely due to the varying stability properties of the states of the organism in mice and humans. In [15], we demonstrate that the dynamic aging signature in humans, characterized by a dynamic frailty index (dFI), exhibits overdamped stochastic fluctuations, typical of dynamically stable system until very late in life (hence the dFI is an adaptive feature in humans). The autocorrelation time of such fluctuations has the meaning of recovery rate (a measure of resilience) and decreases linearly with age.

In [11], we proposed that this linear reduction in resilience results from the interaction (a second-order effect of nonlinear coupling) between dynamic features and the total damage burden (tBA), suggesting that the accumulation of damage registered by (entropic tBA) is the driver of human aging.

We argue that dynamic and entropic aging signatures may be quantified, and hence allow for direct and independent experimental assessment in animal models. Aging is a multidimensional phenomenon, making it difficult to quantify the effects of life-extending interventions using a singular aging clock. Our findings indicate that regression-based clocks uncontrollably mix the dynamic and entropic features with a bias toward capturing the dynamic (reversible) aspects of aging, particularly in mice.

Mice, with their inherent dynamic instability, can exhibit pronounced reactions to life-extension interventions aimed to reduce the dynamic aging signature, which may only yield small (incremental) effects in humans. In [11] and in this letter, we argue that the entropic components of aging may be difficult to reverse. Nonetheless, we expect that future pharmacological advancements that aim to decrease the rate of irreversible damage accumulation could significantly decelerate the aging process. Our understanding is that this is probably the only realistic way to achieve strong life-extension therapies in humans. The identification of both exponential and linear entropic aging patterns in mice not only advances our theoretical grasp of aging in both species but also paves the way for testing of prospective drugs acting on the dynamic and entropic features of aging in short-lived organisms.

## IV. ACKNOWLEDGEMENTS

The authors express their gratitude to Bohan Zhang and Vadim Gladyshev from Harvard Medical School for facilitating access to the data from [19], and to Ferdinand von Meyenn, Stephen Clark, and Wolf Reik from the ETH and Altos labs for providing early access to the data from [12].

## S.I. METHODS

### A. PCA of DNAm data

The data from GSE80672 is a set of site-specific bulk DNAm densities. Due to the limited number of samples representing a very large number of simultaneously measured methylation states, we filter out methylation sites according to their correlation with chronological age (Spearman correlation p-value less than 0.6). In such a way, we selected 1420248 methylation sites (of 1976975 sites in total).

Careful examination of the provided data suggests, that mice there is an independent process, most probably development, that is affecting methylation states at most earlier age. To simplify the analysis below, we filtered samples corresponding to animals younger than 5 months (20 weeks, cf. [9] where a similar consideration was obtained using blood measurements).

To reduce dimensionality of the DNAm measurements by extracting clusters of correlated features in the data, we performed standard scaling and then principal components analysis (PCA).

Let *l*1_*i*_ denote the PC1 loading value of site i. Then 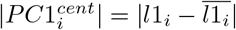.

In the experiment involving Caloric restriction (CR), we fitted the age dependence of DNAm-PCs with the Scipy curve fit function. PC1 age dependency was modelled as 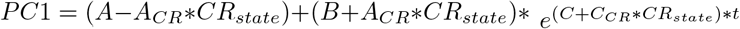. Age was restricted between 6 and 30 months. All the values of the initial parameters were set to 1 except *C* = 0.15. PC2 age dependency was modeled as *PC*2 = *A* + *A*_*CR*_ *CR*_*state*_ + (*B* + *B*_*CR*_ *CR*_*state*_) *t*. Age was not restricted, initial parameters values were all set to 1. Statistical significance of differences in curve parameters were estimated with two-sided t-test. *CR*_*state*_ is a binary variable that equals to 1 if the data point represents a CR mouse and 0 otherwise.

### B. Annotation of DNAm sites and gene set enrichment analysis

Methylation critically influences transcription by modulating transcription factor (TF) binding. Our study aimed to decipher the biological significance of PC1/dFI sites, characterized by |*PC*1^*cent*^ | greater than 0.0015, covering 134,770 sites or the top 10% with the highest PC1 values. We utilized the Gencode vM10 gene annotations (GRCm38/mm10 assembly) for site-to-gene assignments. For each gene, excluding pseudogenes, the promoter region was delineated as 2000 bp upstream and 500 bp downstream of the transcription start site (TSS). Each site was linked to up to two genes based on promoter overlap, discarding sites linked to more than two genes. Genes connected to at least one PC1 site were classified as PC1 genes (4,265 genes), while those linked to at least one PC2 site (|*PC*1^*cent*^| less than 0.001, totaling 1,125,398 sites) were categorized as PC2 (background) genes (18,768 genes). The enrichment analysis was conducted using the EnrichGO function from clusterProfiler version 4.10.0.

### C. Mutual Information (MI) and single-cell methylation analysis

Single-cell DNAm (scDNAm) data, post-preprocessing, consists of binary values indicating the methylation status of specific sites, in addition to a large number of sites without a detected methylation state. Mutual information (MI) serves as a suitable measure to assess the independence between the methylation states of two sites *i* and *j*:

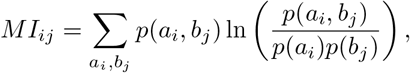

where *p*(*a*_*i*_, *b*_*j*_) denotes the joint probability and *p*(*a*_*i*_) and *p*(*b*_*j*_) the marginal probabilities of methylation states *a*_*i*_ and *b*_*j*_ at sites *i* and *j*, respectively [18].

This study utilized single-cell methylation data, as provided by the authors of [12]. Samples from individuals younger than 5 months and cells that did not meet the methylation quality control (QC) criteria were excluded from the analysis. Detailed information on the retained samples and the number of cells is documented in Supplementary Table 2. In total, 698 scDNAm profiles, aged 8 to 24 months, were retained for analysis. Sites with a methylation level ≤ 30% were categorized as “unmethylated”, and those with a level ≥ 70% as “methylated”. Sites exhibiting intermediate levels of methylation were considered as having no detected methylation state.

For our analysis, we focused on sites identified by the bulk SVD model. Sites with |*PC*1^*cent*^| greater than 0.0015 (top 10%) were classified as PC1 sites, while most of the remaining sites (|*PC*1^*cent*^| less than 0.001) were classified as PC2 sites. The intermediate values |*PC*1^*cent*^| were excluded. Furthermore, we eliminated sites measured in fewer than 50 cells or with a less than 10% representation of the minor state, resulting in 4,810 “PC1” sites and 9,774 “PC2” sites. Pairs of sites with fewer than 10 paired measurements were considered to have non-robust MI estimates and were excluded. We ranked sites by the number of valid MI estimates and selected the top 3,000 sites. From these, we chose 2,000 sites, each with at most 1.7% missing MI values.

To calculate a measure of codependence per DNAm site, we computed the mean MI for each site, considering only sites within the same PC group for the mean MI calculation of each site.

PC1 - MI and PC1 - clock sites association studies were performed via gseapy-1.1.1 prerank function with 100,000 and 1000 permutations, respectively. mHG 1.1 R package was used for the minimal hypergeometric tests.

## Notes

### Competing Interest Statement

The authors have declared no competing interest.

